# Human Pluripotent Stem Cell Derived Organoids Reveal a Role for WNT Signaling in Dorsal-Ventral Patterning of the Hindgut

**DOI:** 10.1101/2024.03.04.583343

**Authors:** Na Qu, Abdelkader Daoud, Daniel O. Kechele, Jorge O. Múnera

## Abstract

The cloaca is a transient structure that forms in the terminal hindgut giving rise to the rectum dorsally and the urogenital sinus ventrally. Similarly, human hindgut cultures derived from human pluripotent stem cells generate human colonic organoids (HCOs) which also contain co-developing urothelial tissue. In this study, our goal was to identify pathways involved in cloacal patterning and apply this to human hindgut cultures. RNA-seq data comparing dorsal versus ventral cloaca in e10.5 mice revealed that WNT signaling was elevated in the ventral versus dorsal cloaca. Inhibition of WNT signaling in hindgut cultures biased their differentiation towards a colorectal fate. WNT activation biased differentiation towards a urothelial fate, giving rise to human urothelial organoids (HUOs). HUOs contained cell types present in human urothelial tissue. Based on our results, we propose a mechanism whereby WNT signaling patterns the ventral cloaca, prior to cloacal septation, to give rise to the urogenital sinus.

## Introduction

The epithelia of the small intestine, colorectum and urothelium are derived from the definitive endoderm. The urothelial epithelium is derived from the ventral portion of the cloaca which forms in the terminal hindgut^1^. The cloaca undergoes septation resulting in the separation of the urogenital sinus (UGS) from the colorectum. Improper septation of the cloaca can result in anorectal malformations (ARMs), a common pathology affecting approximately 1 in 5,000 births^2^. Despite the frequency of ARMs, the mechanisms that pattern the cloaca are not well defined. Understanding these mechanisms would not only provide insight into ARMs but also inform efforts to generate urothelial tissue *in vitro*.

The generation of transplantable urothelial tissue remains an urgent medical need. Patients with bladder cancer^3^, interstitial Cystitis^4^, and bladder exstrophy^5^ often require bladder replacement. The standard of care for all of these patients is construction of neobladder from tissue derived from the patient’s own ileum or sigmoid colon^6^, a procedure which has a high rate (27%) of complications^7^. Expansion of autologous urothelial progenitors could be a source of replacement tissue. However, in the case of bladder cancer, these progenitors may have oncogenic mutations making them unsuitable for transplantation. Therefore, generation of urothelial tissues from human pluripotent stem cells could provide a continuous source of urothelium that bypasses these problems.

We previously developed a method to generate human colonic organoids (HCOs) from human pluripotent stem cells by inducing hindgut patterning. Recently, we demonstrated that by increasing the plating density of the starting cultures, we could generate HCOs with co-developing macrophages. In addition, we also observed urothelium in these cultures^8^. In this study we identified WNT signaling as a pathway that is upregulated in the ventral portion of the cloaca in mice. We found that WNT signaling inhibition prevented urothelial differentiation while WNT signaling activation enhanced urothelial differentiation. Human urothelial organoids (HUOs) generated by activation of WNT signaling contained basal, intermediate, and luminal urothelial cells. Furthermore, HUOs were stratified even in the absence of exogenous factors. Our work reveals that the cloaca is patterned along the dorsal-ventral axis prior to septation and that this patterning is regulated by WNT signaling. Our results provide insights into cloacal septation and anorectal malformations. In addition, HUOs should provide an *in vitro* platform for studying human urothelial development.

## Results

### Genome wide transcriptomic analysis reveals the presence of urothelial-like cell types in HCO cultures

Analysis of RNA-seq data from human intestinal organoids (HIOs) and HCOs (Figure S1A), revealed the presence of urothelial transcripts within HCO cultures but not HIO cultures (Figure S1B). This indicates that the BMP signaling that is required for specification of HCOs, also specified urothelial tissue, which is consistent with the known role of BMP signaling in posterior-ventral development ^9-13^. We confirmed the presence of urothelial tissue expressing KRT13 and found that urothelial organoids developed in similar numbers to colonic organoids (Figure 1A,B; S1C). In addition, we found a small subset of organoids that expressed both urothelial and colonic markers (Figure S1C). These results demonstrate that HCO cultures contain co-developing urothelium.

**Figure 1.**
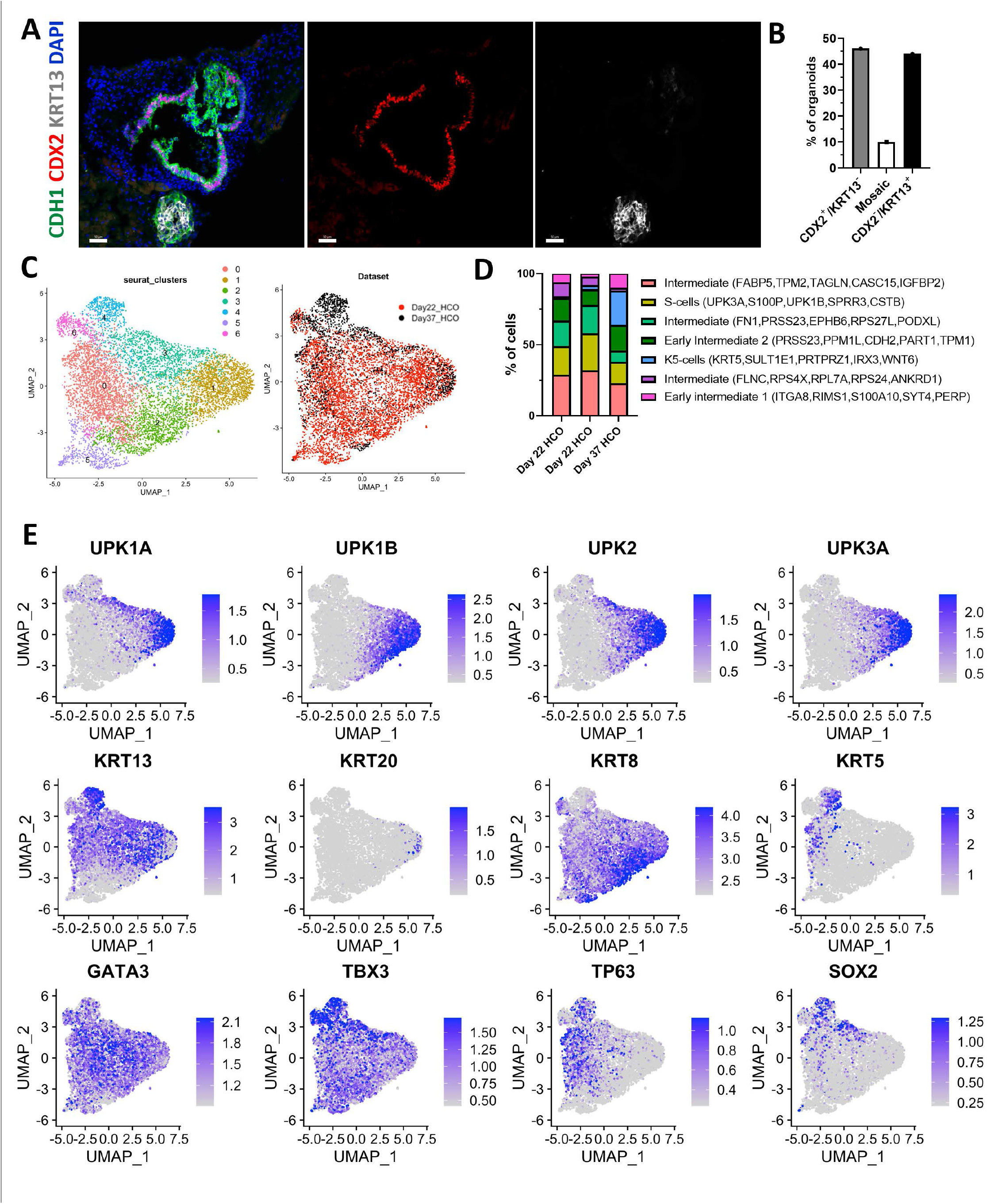
Urothelial tissue is present in HCO differentiations. **(A)** Immunofluorescent staining and **(B)** quantification on day 35 HCO for CDH1 (green), CDX2 (red), and KRT13 (white) and counterstained with DAPI. A total of 41 organoids from 3 separate differentiations were analyzed. **(C)** Uniform manifold approximation and projection (UMAP) Seurat clustering of all urothelial cells from the d22 and d37 HCO cultures. **(D)** Relative proportions of predicted cell types as well as corresponding markers in each HCO sample. **(E)** Feature plots of select urothelial genes including uroplakin genes (*UPK1A, 1B, 2, 3A*), keratin genes (*KRT13, 20, 8 and 5*) and transcription factors (*GATA3, TBX3, TP63, and SOX2*).

To determine the cellular diversity of urothelial cells within 22-day old and 37-day old HCO cultures, we bioinformatically selected the urothelial endoderm from single cell RNA sequencing (scRNA-seq) data on HCO cultures. Uniform manifold approximation and projection (UMAP) analysis of urothelial endoderm revealed the presence of 7 different cell populations within the urothelium including previously described superficial cells (S-cells), and basal keratin 5 expressing cells (K5-cells) ^14,15^ (Figure 1C-D). Although we saw an induction of *KRT14* mRNA in 35-day old HCO cultures in the bulk RNA-seq data (Figure S1B), we did not detect basal KRT14 expressing cells (K14-cells) in the scRNA-seq data (Figure S2A). In addition, we found that the urothelial cells followed a similar developmental trajectory to what occurs in mice since *KRT5* expressing cells were not apparent in 22-day old organoids but present in 37-day old organoids (Figure 1C-E). In summation, these data demonstrate that HCO cultures contain diverse urothelial cell types including urothelial progenitors that undergo maturation and differentiation.

### RNA seq profiling of dorsal and ventral cloaca reveals patterning and increased WNT signaling in the ventral region

To improve the differentiation of HCO cultures towards urothelial tissues, we determined the signaling pathways and transcription factors that are differentially enriched in the ventral versus dorsal cloaca. To achieve this, we analyzed a publicly available dataset from the GenitoUrinary Development Molecular Anatomy Project (GUDMAP). In this dataset, the ventral and dorsal cloaca from e10.5 mouse embryos were separated using laser capture microdissection, and RNA was collected from the samples and subjected to RNA-seq analysis (Figure 2A). Principal component analysis revealed that ventral and dorsal cloaca samples clustered into distinct groups based on their transcriptomic profiles (Figure 2B). We then performed gene ontology analysis on transcripts that were enriched in ventral versus dorsal cloaca. This analysis revealed a strong enrichment for WNT signaling in the ventral cloaca (Figure 2C) indicating that there is a gradient of WNT signaling along the dorsal-ventral axis in the cloaca. Gene ontology analysis of dorsal cloaca revealed a strong enrichment for epithelial development and embryonic morphogenesis (Figure S3A).

**Figure 2.**
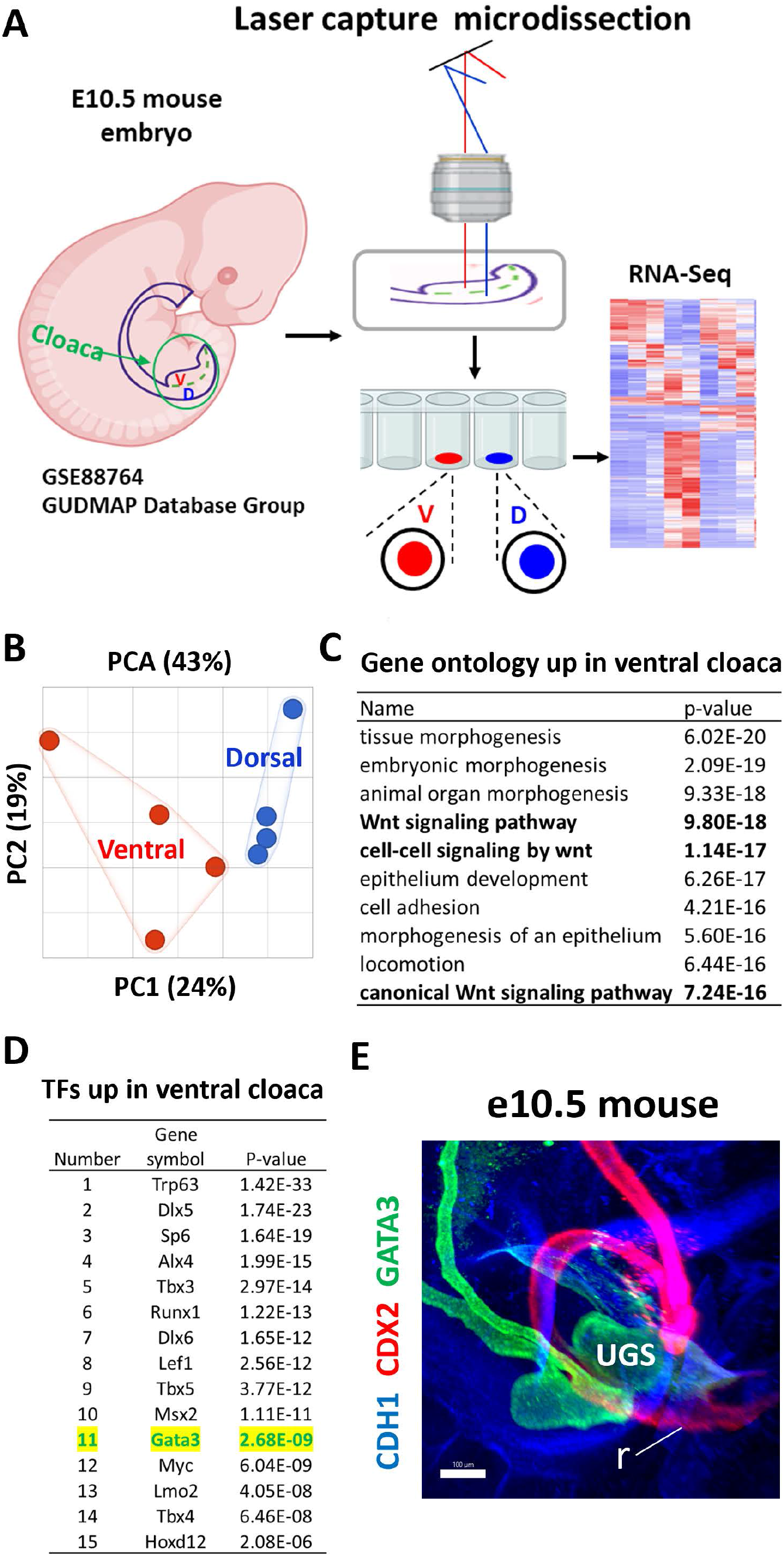
WNT signaling is increased in ventral versus dorsal cloaca. **(A)** Schematic of cloacal tissue isolation used to generate RNA-seq data for GSE88764. **(B)** Principal component analysis of samples from dorsal cloaca versus ventral cloaca. 4 biological replicates per condition were analyzed. **(C)** Gene ontology analysis of genes that are upregulated in the ventral versus dorsal cloaca. **(D)** Table of most upregulated transcription factors in ventral cloaca. **(E)** Whole-mount immunofluorescence staining of an e10.5 embryo stained for CDH1 (blue), the pan-intestinal marker CDX2 (red) and urothelial marker GATA3 (green).

To examine the transcription factors that are enriched in the ventral cloaca, we filtered out the transcription factors present in the differentially expressed genes. We observed enrichment of several transcripts known to be enriched in bladder tissue including *Trp63*^16,17^ and *Gata3*^18^ (Figure 2D). Dorsal cloaca was enriched in colon transcription factors including *Cdx2*^19,20^ and the intestinal mesenchyme marker *Nkx2-3*^21,22^ (Figure S3B). Wholemount immunofluorescence staining of e10.5 mouse embryos, when the colorectum and UGS begin to septate, revealed that GATA3 is expressed in the UGS portion of the cloaca, and CDX2 is expressed in the colorectal portion of the cloaca with no overlap between these two factors (Figure 2E, Figure S2B). Our immunostaining for GATA3 also stained other tissues which are known to express GATA3 including the nephric ducts (which are derived from mesoderm), consistent with previous studies^23-26^, and confirming the specificity of our staining. Immunostaining for Runx1, which was also enriched in the ventral cloaca, revealed that RUNX1 marks the most ventral portion of the cloaca (Figure S2C). Our findings reveal that WNT signaling is elevated in the ventral cloaca compared to the dorsal counterpart and that CDX2, GATA3 and RUNX1 mark the dorsal and ventral cloaca prior to septation.

### WNT activation in human hindgut cultures induces urothelial specification

Based on the data from mice, we determined if WNT signaling could pattern HCO cultures along the dorsal-ventral axis. To do this, we patterned organoids using BMP but also treated, either with an inhibitor of catenin related transcription 3 (ICRT3), or the GSK3B inhibitor CHIR99021 (Figure 3A). Principal component analysis of the RNA-seq data revealed that WNT inhibited and WNT activated samples clustered into distinct groups based on their transcriptomic profiles (Figure 3B). Gene ontology analysis on transcripts that were enriched in WNT activated organoids, revealed an enrichment for proliferation associated genes and a significant enrichment for WNT signaling and urogenital system development (Figure 3C). WNT inhibited organoids were enriched in circulatory system development and tube development (Figure S3C). Analysis of transcription factors revealed that WNT inhibition resulted in the increased expression of several factors known to be involved in colorectal development , such as HLX^27^, or expressed specifically in intestinal tissue such as ISX^28^. (Figure S3D). In contrast, WNT activation resulted in the increased mRNA expression of several factors known to be involved in urothelial development including GRHL3^29^ and TP63^16^ as well as the ventral cloaca marker GATA3 (Figure 3D). Furthermore, 6 of the top 12 transcription factors that were upregulated in WNT activated day 10 hindgut cultures (Figure 3D), were also upregulated in the ventral cloaca including *ALX4, DLX5, DLX6, LEF1, MSX2* and *TP63*. These results indicate that WNT signaling dictates the lineage decision between urothelial and colorectal progenitors in human hindgut cultures allowing the biased generation of human urothelial organoids (HUOs).

**Figure 3.**
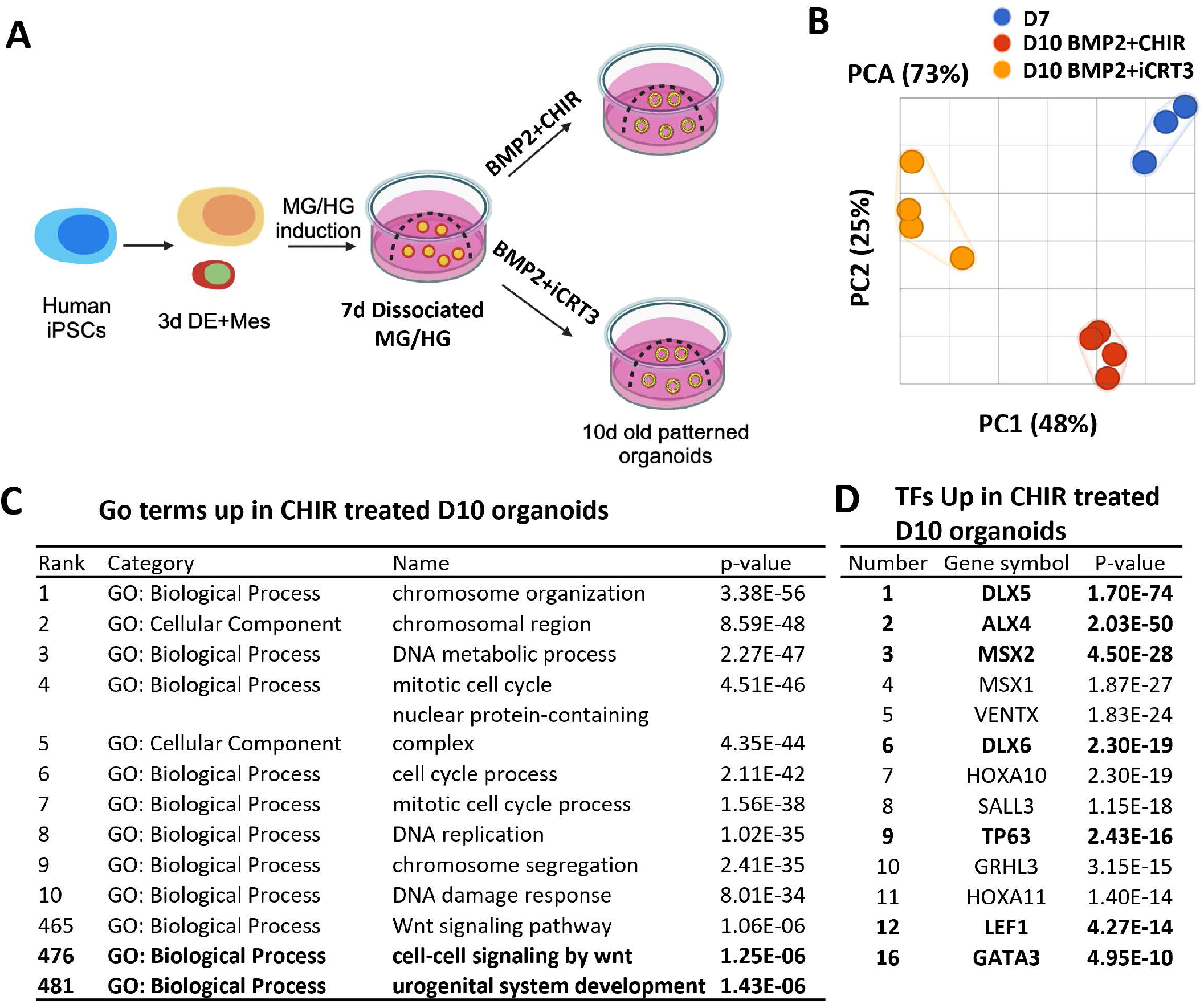
WNT signaling regulates lineage fate of hindgut cultures. **(A)** Schematic of WNT manipulation during patterning of human hindgut cultures. **(B)** Principal component analysis comparing day 7 midgut/hindgut cultures, day 10 BMP + ICRT3, and day 10 BMP + CHIR cultures. Samples were from 3-4 separate differentiations. **(C)** Gene ontology analysis of genes upregulated in day 10 BMP + CHIR cultures compared to day 10 BMP + ICRT3 cultures. (D) Table of most upregulated transcription factors in day 10 BMP + CHIR cultures compared to day 10 BMP + ICRT3 cultures. Transcription factors in (D) that overlap with those upregulated in ventral cloaca are shown in bold.

### WNT activated organoids maintain ventral identity and generate urothelial cell types

To determine if WNT manipulations resulted in stable specification of colonic or urothelial tissue, we grew organoids for 35 days (Figure 4A). Quantification of organoids immunostained with CDX2 to mark colonic epithelial cells and GATA3 to mark urothelial endoderm, revealed that ICRT3 treatment resulted in differentiation towards CDX2+ organoids. In contrast, activation of WNT signaling resulted in differentiation towards GATA3+ urothelial organoids (Figure 4B). To determine the global transcriptional changes associated with WNT manipulation, we performed bulk RNA-seq on 35-day old organoids. Principal component analysis (PCA) revealed that BMP2+ICRT3 treated cultures clustered separately from BMP2+CHIR treated cultures (Figure 4C). To broadly interrogate the tissue identity of BMP2+ICRT3 and BMP2+CHIR treated cultures we compared them with published data sets of human fetal large intestine and human fetal bladder. Principal component analysis revealed that primary tissues isolated from fetal bladder and our organoids clustered together along principal component 1 (PC1) axis, which accounted for 56% of the cumulative variation among samples, reflecting the persistence of minor population (11%) of HUOs in BMP2+ICRT3 treated cultures (Figure 4D). The second principal component (PC2) which accounts for 14% of cumulative variation, revealed that BMP2+ICRT3 treated cultures clustered closer to human fetal large intestine and BMP2+CHIR treated cultures clustering closer to human fetal bladder (Figure 4D). Examination of genes that were upregulated in response to WNT inhibition revealed that BMP2+ICRT3 treated cultures were enriched in the fetal large intestinal transcriptomic signature which included the expression of the pan-intestinal transcription factors *CDX1* and *CDX2*^20^, as well as the colon specific transcription factor FXYD2^30^ (Figure S4A). In contrast, BMP2+CHIR treated cultures were enriched in a fetal bladder transcriptional signature which included the transcription factors *GATA3*^*31*,*32*^, *GRHL3*^29^ and *TP63*^16^ (Figure S4B). These results confirm that BMP2+ICRT3 treatment stably biases hindgut cultures toward large intestine and that BMP2+CHIR treatment stably biases the differentiation of hindgut cultures into urothelium.

**Figure 4.**
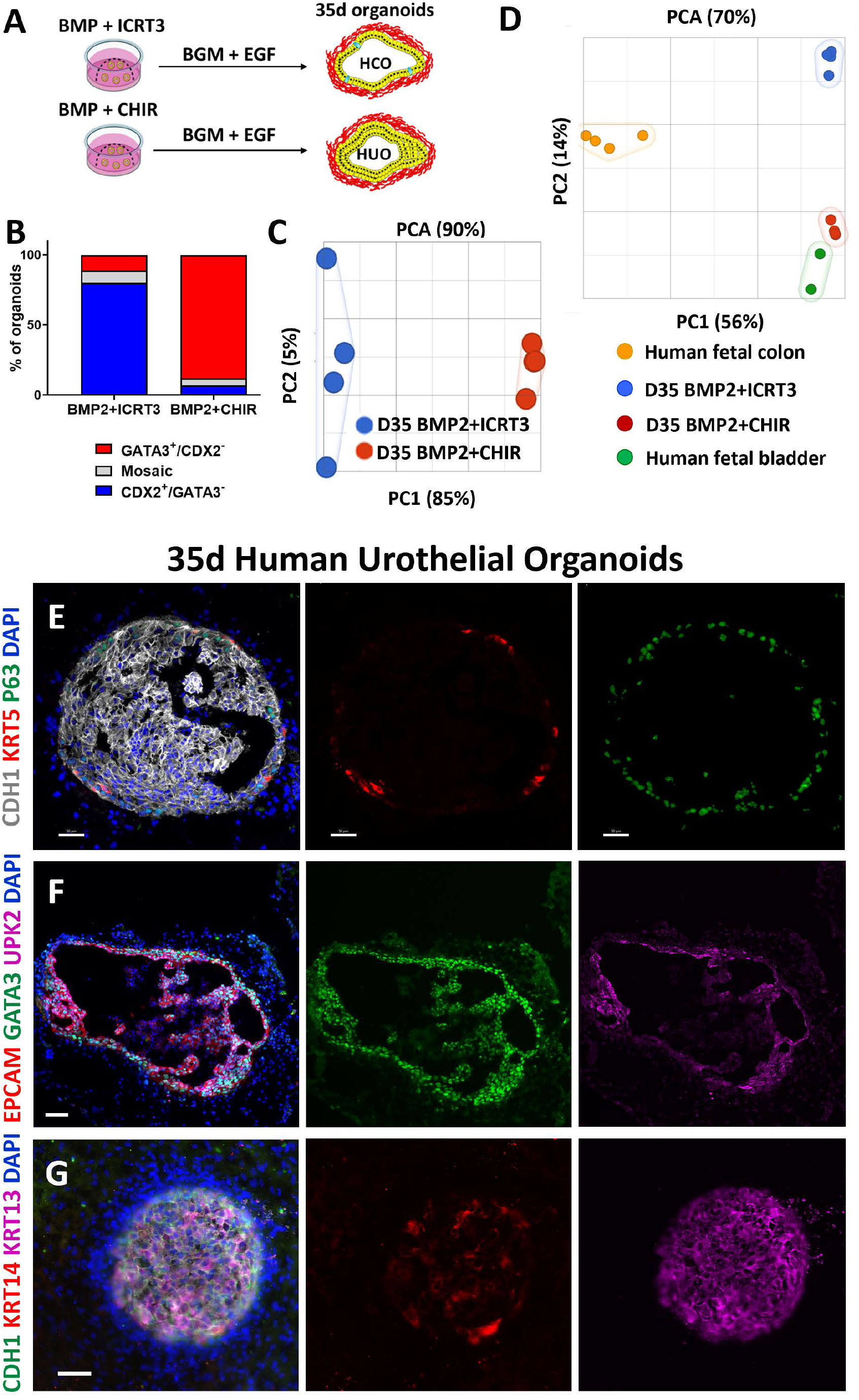
WNT patterned hindgut cultures generate urothelial tissue. **(A)** Schematic of long-term growth of BMP + CHIR cultures and BMP + ICRT3 cultures to 35 days. **(B)** Quantification of organoids stained for CDX2 and GATA3. **(C)** Principal component analysis comparing day 35 BMP + ICRT3, and day 35 BMP + CHIR cultures. Samples from 4 separate differentiations are shown. **(D)** Principal component analysis comparing two day 35 cultures, human fetal colon, and human fetal bladder. Immunofluorescence staining of day 35 BMP + CHIR (HUO) cultures stained for **(E)** CDH1 (white), KRT5 (red), TP63 (green), **(F)** EPCAM (red), GATA3 (green), UPK2 (magenta), and **(G)** CDH1 (green), KRT14 (red), KRT13 (magenta). **(E-G)** Samples were counterstained with DAPI. Images are representative of a minimum of 5 organoids per differentiation from 4 separate differentiations.

We next used hypergeometric means test to determine the probability that BMP2+ICRT3 and BMP2+CHIR organoids share similar patterns of tissue-specific gene expression with human fetal large intestine and bladder respectively (Figure S4C). A total of 1,875 transcripts are expressed in the human fetal large intestine and in BMP2+ICRT3 treated organoids compared to human fetal bladder or BMP2+CHIR treated organoids, a proportion that is exceedingly unlikely by chance alone (P = 2.9 x 10^-299^). Conversely, the set of genes that is up-regulated in BMP2+CHIR treated organoids is highly enriched for genes that are up-regulated in the human fetal bladder compared to human fetal large intestines. This analysis concluded that BMP2+ICRT3 treated organoids are most similar to human fetal large intestine and BMP2+CHIR treated organoids are most similar to human fetal bladder. Taken together, these data suggest that we have developed a robust method to differentiate PSCs into human urothelial tissue which we termed HUOs.

To confirm the presence of urothelial cell types in HUO (BMP2+CHIR) cultures, we examined the expression of other urothelial markers by immunofluorescence staining in 35-day old organoids. HUOs expressed the basal marker KRT5 as well as TP63 (Figure 4E). Furthermore, HUOs expressed GATA3 and UPK2 indicating that organoids were urothelial and that they generated umbrella cells (Figure 4F). In addition, HUOs expressed the urothelium enriched marker KRT13 and the basal marker KRT14 (Figure 4G). HUOs were also multilayered indicating that epithelial stratification occurs spontaneously in our cultures. Collectively these results indicate that WNT activation biases the differentiation of hindgut cultures into HUOs that undergo stereotypical urothelial differentiation and contain basal K5-cells, S-cells and intermediate cells.

## Discussion

Lineage tracing studies have demonstrated that the colorectum, bladder and prostate all develop from common hindgut progenitors^17^. Therefore, the presence of urothelial organoids in HCO cultures reflects the common origin of colorectal and urothelial tissues. Multiple studies have reported the generation of urothelium from human pluripotent stem cells^33-35^, although only one of these protocols generated stratified urothelium^33^. However, generation of stratified urothelium required the growth of urothelial cells on transwells with the addition of the combination of the PPAR-γ agonist troglitazone, the EGFR inhibitor PD153035, and FGF10. In contrast, our HUOs generated stratified urothelial tissue without the addition of these factors. This indicates that HUOs are a suitable model for examining basal to luminal differentiation of urothelial progenitors and that HUO cultures endogenously supply the factors that are required for stratification of the urothelium.

Our results also demonstrate the presence of KRT5 expressing cells in our human hindgut cultures. In both murine models and in human pluripotent stem cell cultures, the origin of basal cells in the urothelium remains unknown. In mice, the appearance of basal cell populations coincides with increased urinary outflow in mouse embryos^36^ and it has been suggested that this stress stimulus might regulate the appearance or expansion of these cells^37^. Urothelium within our human hindgut cultures display maturation and KRT5 expressing cells are only present in 37-day old organoids. Our findings suggest that the appearance of these cells is intrinsically regulated in our organoid cultures. However, urinary flow may enhance the numbers of KRT5 expressing cells that appear. Examination of multiple timepoints between day 22 and day 37 should allow us to elucidate the transcriptional programs that are involved in the generation of KRT5 expressing basal cells.

We found that WNT signaling is enriched in the ventral cloaca. Multiple studies have implicated WNT signaling in urothelial development^38-40^. Interestingly, a previous GWAS study identified duplications in the WNT inhibitor *DKK4* in patients with ARMs^41^. Our results indicate that WNT signaling biases the differentiation of hindgut progenitors toward urothelial fate. Therefore, CDX2/GATA3 co-staining could reveal if ARMs caused by WNT signaling modulation are a result of expanded ventral or dorsal cloacal progenitors. For instance, we would predict that patients with mutations in DKK4 would exhibit expansion of the GATA3 domain into the dorsal portion of the cloaca. Interestingly, loss of CDX2, and ectopic expression of GATA3 were reported in the rectum of a fetus with urorectal septum malformation sequence (URSM)^42^ suggesting that the maintenance of the CDX2 and GATA3 boundary may be important for proper septation.

The septation of the cloaca into UGS and colorectum resembles the septation of the esophagus and trachea from a common foregut tube. Consistent with this observation, genetically engineered mice with perturbed Sonic Hedgehog^43^, or WNT^44^ signaling have defects in septation of the esophagus and trachea as well as defects in the septation of the cloaca into UGS and colorectum^38,45^. Furthermore, tracheoesophageal fistula and anorectal malformations occur in conjunction with one another in human VATER syndrome (<u>V</u>ertebral, <u>A</u>norectal, <u>T</u>racheo<u>e</u>sophageal, <u>R</u>enal)^46^. Septation of the esophagus and trachea in mice requires the dorsoventral patterning and specification of the common foregut tube into a dorsal Sox2 expressing esophageal domain and a ventral Nkx2.1 expressing tracheal domain^47^. Our results suggest that the cloaca may be patterned along the dorsal-ventral axis in a similar manner, with WNT signaling inducing the GATA3 expression domain and inhibiting the CDX2 expression domain of the cloaca.

### Limitations of Study

We found that WNT signaling is enriched in the ventral cloaca of mice. We then tested whether WNT signaling can pattern human hindgut cultures by using small molecule inhibitors and activators of the pathway. Although RNA-seq data confirmed that WNT signaling was modulated by the small molecules used, these small molecules can have off target effects that may contribute to the observed phenotypes. Future gain of function studies with constitutively active Beta-catenin and loss of function studies with hPSCs where Beta-catenin is targeted using CRISPR-Cas9 should directly address these limitations. In addition, establishment of an orthotopic transplantation model should determine the utility of HUOs for regenerative medicine.

## Acknowledgements

We would like to thank Cassie M. Cleary, Kaleigh Doyle, Avery C. Jackson for their assistance in experiments. We would like to thank the research cores at MUSC including the Cell Models Core and the Advanced Imaging Core at MUSC. This research was supported by grants from the NIH. This work was supported by NIDDK 1R56DK129575, from the MUSC COBRE CDLD (P20 GM130457) and DDRCC (P30 DK123704).

## Contributions

J.O.M. conceived the study and experimental design, N.Q. performed and analyzed experiments and co-wrote the manuscript. A.D, performed experiments. D.O.K performed data analysis. All authors contributed to the editing of the manuscript.

## Declaration of interests

No competing interests declared related to this work.

**Figure S1 related to figure 1. Urothelial markers are present in HCO differentiations. (A)** Schematic of HIO versus HCO differentiation. **(B)** Heatmap of urothelial genes in HIOs compared to HCOs. Heatmap is based on TPM values from RNA-seq data. **(C)** Examples of immunostaining of colon organoids, urothelial organoids, mosaic organoids based on staining for CDH1 (green), CDX2 (red), and KRT13 (white) and counterstained with DAPI .

**Figure S2 related to figure 1. Additional marker(s) expressed in mouse ventral hindgut and urothelium from HCO differentiations. (A)** Feature plots of additional urothelial genes. Wholemount immunofluorescence staining of day an e10.5 mouse embryos for **(B)** CDH1 (blue), CDX2 (red), GATA3 (green), and **(C)** FOXA2 (red), RUNX1 (green).

**Figure S3 related to figure 2 and 3. Gene ontology and transcription factor expression in WNT modulated 10-day hindgut organoids. (A)** Gene ontology analysis of genes that are upregulated in the dorsal versus ventral cloaca. **(B)** Table of most upregulated transcription factors in dorsal cloaca. **(C)** Gene ontology analysis of genes upregulated in day 10 BMP + ICRT3 cultures compared to day 10 BMP + CHIR cultures. **(D)** Table of most upregulated transcription factors in day 10 BMP + ICRT3 cultures compared to day 10 BMP + CHIR cultures. TFs that are common in (**C**,**D)** are shown in bold.

**Figure S4 related to figure 4. Dorsal ventral patterning by WNT modulation is stable. (A)** Heatmap of human fetal large intestinal transcripts and their expression in BMP + ICRT3 cultures compared to day 35 BMP + CHIR cultures. **(B)** Heatmap of human fetal bladder transcripts and their expression in day 35 BMP + CHIR cultures compared to day 35 BMP + ICRT3 cultures. **(C)** Hypergeometric means tests comparing thetranscriptomes of human fetal large intestine with HCOs and human fetal bladder with HUOs.

## Methods

### Resource Availability

#### Lead Contact

For additional information and requests for resources, reagents, or codes, please contact Jorge Munera

## Materials Availability

All human induced pluripotent stem cells lines are available upon request with appropriate MTA and are listed in Key Resource Table.

## Data and Code Availability

- Some RNA-seq and scRNA-seq datasets analyzed in this study are publicly available in the GEO repository with accession number GSE240363. All accession numbers used in this study are listed in the Key Resource Table.
- RNA-seq data of e10.5 mouse dorsal and ventral cloaca is publicly available in the GEO repository with accession number GSE88764.
- Bulk RNA-seq datasets for day 10 and 35 WNT inhibition and WNT activation will be deposited into GEO.
- All scRNA-seq analysis was performed in R using previously generated codes, which are referenced in the Key Resource Table. This paper does not report original code.
- Additional information, original images, or scripts needed for further analysis are available upon request.

### Experimental Models

#### Human PSC Line Generation and Maintenance

The human ESC (H1, WiCell, RRID: CVCL_9771) and control iPSC line 72.3 (RRID: CVCL_A1BW) were obtained from the Cincinnati Children’s Hospital Medical Center (CCHMC) Pluripotent Stem Cell Facility. Human iPSC 72.3 were generated from foreskin fibroblasts using episomal plasmids. All lines were derived from male donors. HPSCs were maintained in mTeSR1 (StemCell Technologies) on hESC-qualified Matrigel (Corning) coated Nunclon Delta Surface 6-well plates (Thermo Scientific). Spontaneous differentiation was manually removed and cells were passaged every 3-4 days using Dispase. Cells were routinely confirmed to be karyotypically normal and mycoplasma negative.

### Method Details

#### Directed Differentiation of human PSCs into HIO, HCOs and HUOs

Generation of HIOs and HCOs has been previously published ^48,49^. Briefly, human ES and iPSCs were plated as single cells in mTeSR1 media plus ROCK inhibitor Y27632 (10 μM; Stemgent) on hESC-qualified Matrigel (Corning)-coated 24-well plate at 150,000 cells per well. Beginning the next day, cells were treated with Activin A (100 ng/ml; Cell Guidance Systems) for three days in RPMI 1640 (Invitrogen) containing increasing concentrations of 0%, 0.2%, and 2.0% define fetal bovine serum (dFBS; Invitrogen). For endoderm induction of iPSC72.3, BMP4 (15 ng/ml) was added to the first day of Activin A treatment. Following definitive endoderm induction, cells were treated for 4 days with FGF4 (500 ng/ml; R&D Systems) and CHIR99021 (3 μM; Stemgent) in RPMI 1640 with 2.0% dFBS to generate 3-dimensional mid-hindgut spheroids. For WNT inhibition and activation studies we used mid/hindgut monolayer that was dissociated into clumps as previously described^50^. These mid/hindgut clumps were embedded in basement-membrane Matrigel (BD Biosciences) and subsequently grown in Advanced DMEM/F12 (Invitrogen) supplemented with N2 (Invitrogen), B27 (Invitrogen), L-glutamine, 10 μM HEPES, penicillin/streptomycin, and EGF (100 ng/ml; R&D Systems). For proximal HIO and HCO specification, Noggin (100 ng/ml; R&D Systems) or BMP2 (100 ng/ml; R&D Systems) was added for the first three days of three-dimensional growth, respectively. For WNT modulation experiments, BMP2 (100 ng/ml; R&D Systems) and either ICRT3 (50μM; Selleckchem) or CHIR99021 (3 μM; Stemgent) was added for the first three days of three-dimensional growth. Organoids were transferred to new Matrigel following 2-3 weeks.

### Tissue processing, immunofluorescence, and microscopy

Tissues were fixed for 0.5-1 hr in 4% paraformaldehyde (PFA) on ice. Organoids frozen in OCT. OCT sections were blocked using donkey serum (5% serum in 1× PBS plus 0.5% Triton-X) for 30 min and incubated with primary antibody overnight at 4 °C. Slides were then washed 3X with 1X PBS plus 0.5% Triton-X and incubated in secondary antibody with DAPI in blocking buffer for 2 hr at room temperature (23°C). Please see Table 1 for list of antibodies. Slides were then washed 2X with 1X PBS plus 0.5% Triton-X followed by a final wash in 1X PBS. Slides were then mounted using Fluoromount-G® (SouthernBiotech). Images were captured on a Nikon A1 confocal microscope (CCHMC) or a Zeiss LSM 880 NLO confocal microscope (MUSC) and analyzed using NIS Elements (Nikon) or Imaris Imaging Software (Bitplane).

For wholemount staining of mouse embryos, fixed embryos were washed with PBS and then permeabilized in PBST overnight at 4 °C on a rocking platform. Embryos were then blocked for 6-8 hours on a rocking platform, incubated in primary antibody overnight at 4 °C on a rocking platform, washed 5X with PBST, incubated in secondary antibody overnight at 4 °C on a rocking platform, washed 2X with PBST followed by a final wash in PBS. Embryos were then dehydrated by washing 3 times in 100% Methanol and cleared using Murray’s clear (2 parts benzyl benzoate and 1 part benzyl alcohol).

Images were captured on a Nikon A1 confocal microscope using Z-correction and analyzed using Imaris Imaging Software (Bitplane).

### RNA isolation and Bulk RNA-seq sequence assembly abundance estimation

RNA was extracted using NucleoSpin® RNA extraction kit (Macharey-Nagel) according to manufacturer’s protocols. RNA library construction and RNA sequencing was performed by BGI using the DNBseq platform. QC analysis was performed with data from the DNBseq platform and downstream analysis was performed using Partek® Flow® software. Heatmaps were generated using TPM tables that were then converted into heatmaps using Morpheus (Broad Institute). H1 cell derived HIO and HCO data can be found by accession number GSE240363 . WNT inhibited or activated hindgut organoids are also from three independent differentiations done using the IPSC72.3 line. Human fetal RNA-seq data were downloaded from GEO. Human fetal colon data can be found via the accession number GSE18927. Human fetal bladder data can be found via the accession number GSE78563.

### Single Cell RNA-seq Analysis

We used previously published scRNA-seq data which can be found by accession number GSE240363, for our analysis. Raw scRNA-seq data was converted to FASTQ files and then aligned to the human genome [hg19 d37 HCO or hg38 for d22 HCO] using CellRanger v3.0.2 (10x Genomics). Individual analysis, including quality controls and clustering, of all datasets was first performed using Seurat [v3.2.3] ^51^ in R [v3.6.3]. Basic filtering parameters for gene detection included greater than or equal to 3 cells and all cells with minimum 100 detected genes and maximum 7,500 genes. Percent.mito parameter was all cells less than 20. Data was normalized using SCTransform in Seurat. Cell cycle effect was regressed out using previously established methods in Seurat. Normalized expression levels underwent principal component analysis (PCA) followed by uniform manifold approximation and projection (UMAP) ^52^ with subsequent Louvain clustering. Cluster resolution was changed to increase or decrease number of clusters identified. Datasets from two independent d22 HCO replicates were merged using Seurat prior to normalization, PCA, UMAP, and clustering. Marker genes were determined using ‘FindAllMarkers’ function (Wilcoxon rank-sum test). Clusters were annotated manually using unbiased methods.

### Data Quantification

For experiments involving patterned mid/hindgut clumps and *in vitro* grown organoids, “n” represents the number of biological replicates (2-3 wells were collected for each replicate). Graphs were generated in GraphPad Prism. All figures were generated using Adobe Photoshop or Illustrator and model schematics were animated using BioRender.

### Declaration of generative AI and AI-assisted technologies in the writing process

During the preparation of this work the author(s) used ChatGPT to improve language and readability. After using this tool/service, the author(s) reviewed and edited the content as needed and take(s) full responsibility for the content of the publication.

