## Supplementary materials for "Human Pluripotent Stem Cell Derived Organoids Reveal a Role for WNT Signaling in Dorsal-Ventral Patterning of the Hindgut"

**Figure S1 related to figure 1. Urothelial markers are present in HCO differentiations.**

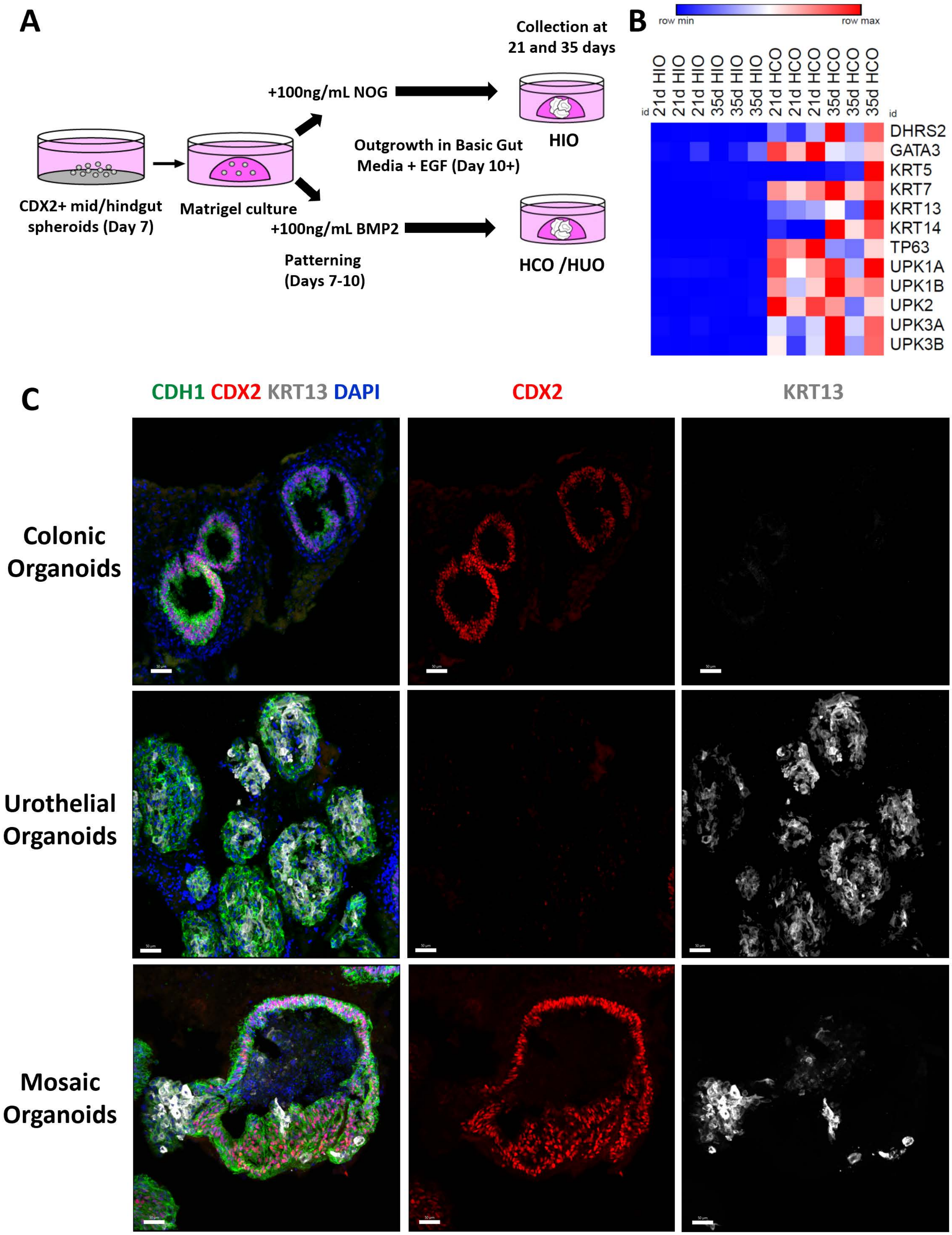

**Figure S2 related to figures 1 and 2. Additional markers expressed in urothelium from HCO cultures and in ventral cloaca of mouse embryos.**

**A**

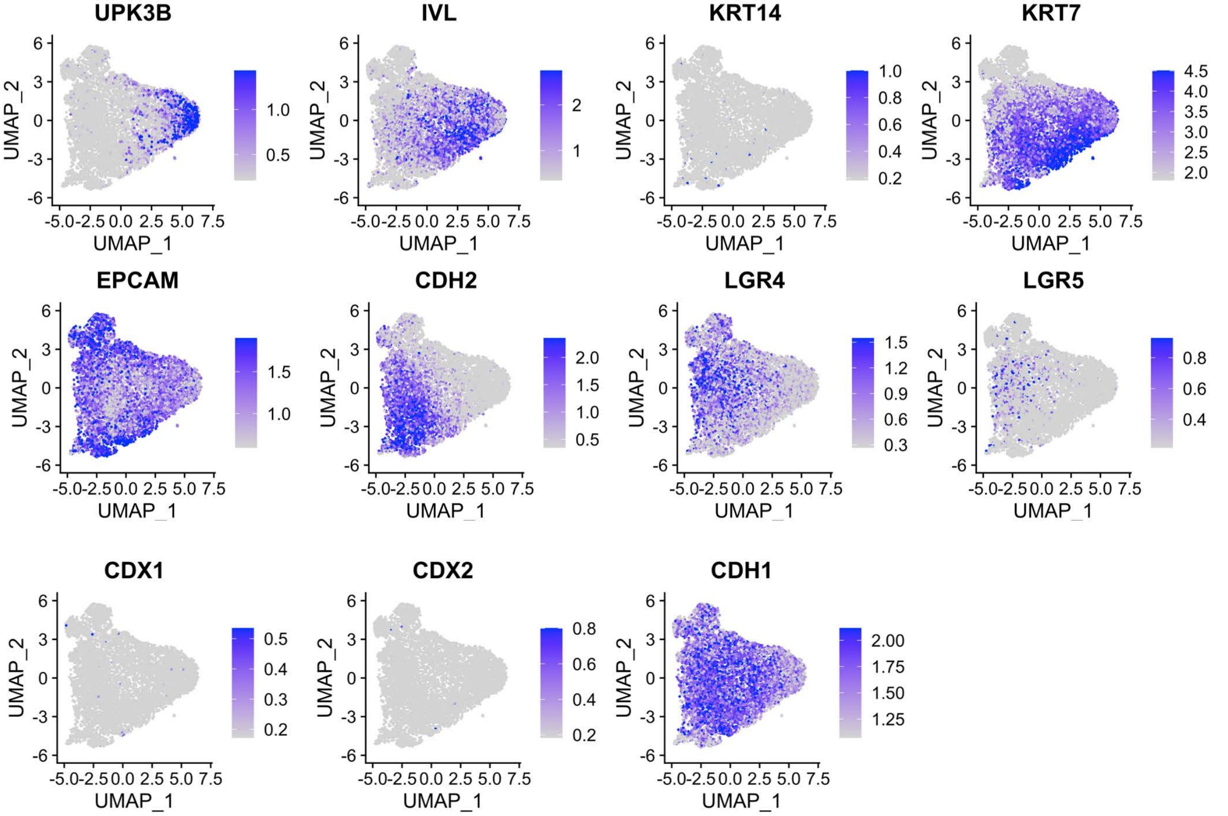


**B** **e10.5 mouse embryos.**

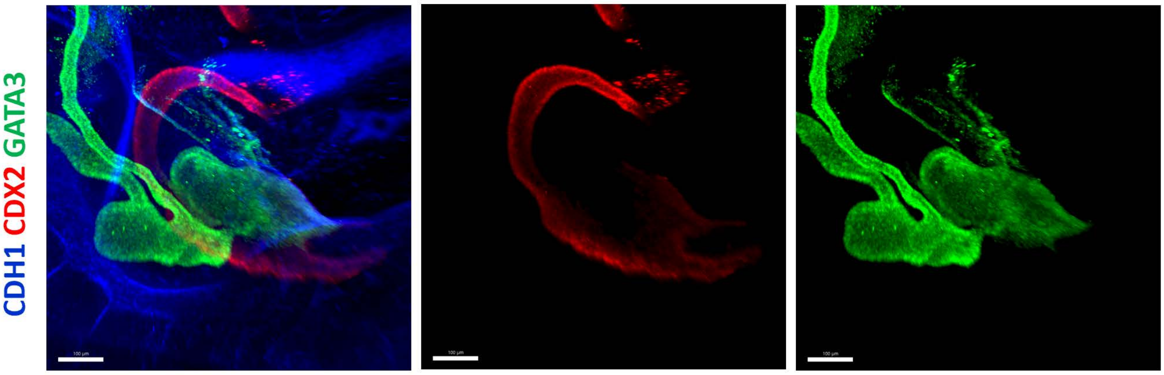


**C**

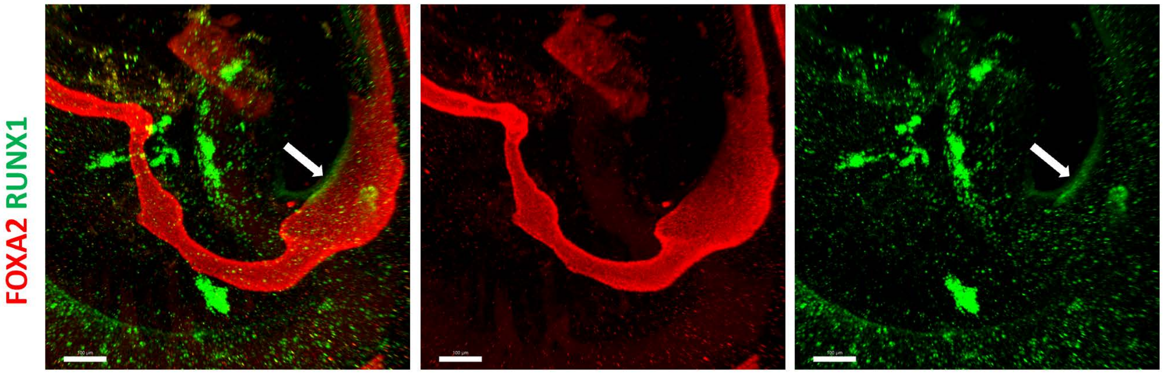

### Figure S3 related to figures 2 and 3.

## A

##### Up in dorsal region of cloaca

| Category | Name | p-value |
|---|---|---|
| GO: Biological Process | epithelium development | 3.30E-10 |
| GO: Biological Process | embryonic morphogenesis | 3.37E-09 |
| GO: Biological Process | head development | 3.54E-09 |
| GO: Biological Process | brain development | 4.17E-09 |
| GO: Biological Process | tube development | 7.81E-09 |
| GO: Biological Process | tissue morphogenesis | 9.04E-09 |
| GO: Biological Process | morphogenesis of an epithelium | 1.26E-08 |
| GO: Biological Process | epithelial cell development | 2.86E-08 |
| GO: Biological Process | central nervous system development | 3.09E-08 |
| GO: Biological Process | response to decreased oxygen levels | 6.26E-08 |

## C

##### Up in ICRT3 treated D10 organoids

| Category | Name | p-value |
|---|---|---|
| GO: Biological Process | circulatory system development | 3.89E-39 |
| GO: Biological Process | tube development | 7.20E-34 |
| GO: Biological Process | tube morphogenesis | 6.90E-33 |
| GO: Biological Process | regulation of anatomical structure morphogenesis | 1.23E-31 |
| GO: Biological Process | cell adhesion | 1.81E-30 |
| GO: Biological Process | cell morphogenesis | 1.94E-30 |
| GO: Biological Process | vasculature development | 2.50E-29 |
| GO: Biological Process | blood vessel development | 1.11E-28 |
| GO: Biological Process | anatomical structure formation involved in morphogenesis | 5.99E-28 |
| GO: Biological Process | vesicle-mediated transport | 8.70E-28 |

## B

##### TFs up in dorsal region of cloaca

| Number | Gene symbol | P-value |
|---|---|---|
| 1 | Ppargc1a | 2.03E-09 |
| **2** | **Hhex** | **1.15E-08** |
| 3 | Cdx2 | 2.89E-07 |
| 4 | Fmnl2 | 8.21E-06 |
| 5 | Hand1 | 1.16E-05 |
| 6 | Lbh | 2.28E-05 |
| **7** | **Foxa1** | **6.32E-05** |
| 8 | Nkx2-3 | 1.30E-04 |
| 9 | Hlf | 1.74E-04 |
| 10 | Mcm4 | 2.04E-04 |
| 11 | Jun | 2.25E-04 |
| 12 | Vgll2 | 2.37E-04 |
| 13 | Enpp2 | 5.29E-04 |
| 14 | Plagl1 | 7.70E-04 |
| 15 | Pdlim1 | 7.83E-04 |

## D

##### TFs up in ICRT3 treated D10 organoids

| Number | Gene symbol | P-value |
|---|---|---|
| 1 | ONECUT2 | 1.51E-79 |
| 2 | CEBPA | 4.62E-45 |
| 3 | HLX | 3.70E-41 |
| 4 | ISX | 1.16E-40 |
| 5 | RFX6 | 2.69E-39 |
| 6 | HOXA1 | 1.77E-29 |
| 7 | TFCP2L1 | 1.39E-25 |
| **8** | **FOXA1** | **5.44E-24** |
| 9 | HNF1B | 6.26E-21 |
| 10 | RARA | 1.90E-20 |
| 11 | IRF8 | 2.67E-20 |
| 12 | HOXA3 | 3.19E-19 |
| 13 | NR2F2 | 2.46E-16 |
| **14** | **HHEX** | **7.91E-16** |
| 15 | NR5A2 | 1.14E-15 |
| 16 | JDP2 | 2.40E-15 |
| 17 | GATA6 | 1.25E-14 |
| 18 | BCL6 | 1.89E-14 |
| 19 | KLF2 | 3.39E-14 |
| 20 | NR2F1 | 7.36E-12 |

**Figure S4. Dorsal ventral patterning by WNT modulation is stable.**

**A** **Human fetal colon signature**

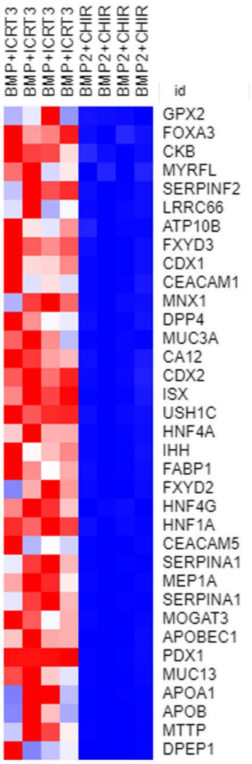


**B** **Human fetal bladder signature**

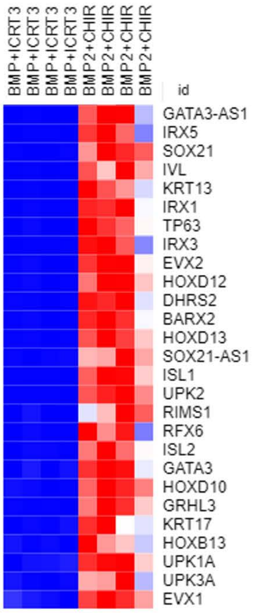


**C** **Up in BMP+ICRT3 vs BMP+CHIR** **Up in HFC vs HFB**

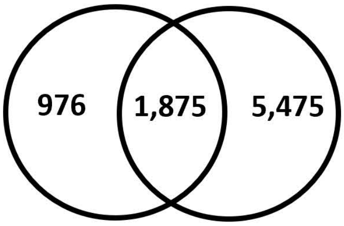


**p < 2.898e-299**

**D** **Up in BMP+CHIR vs BMP+ICRT3** **Up in HFB vs HFC**

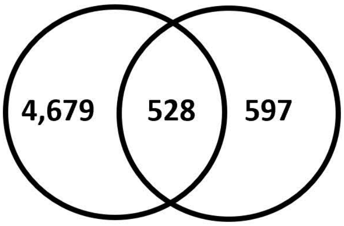


**p < 4.832e-65**

| Antigen | Species | Cat# | RRID |
| --- | --- | --- | --- |
| CDH1 | goat | AF648 | RRID:AB_355504 |
| CDH1 | rat | MAB7481 | RRID:AB_2076679 |
| CDX2 | mouse | MU392A5UC/NC1083496 | RRID:AB_2650531 |
| CDX2 | rabbit | 235R-14 | RRID:AB_1516797 |
| EPCAM | mouse | 248M-94 | RRID:AB_1516851 |
| FOXA2 | goat | sc-6554 | AB_2262810 |
| GATA3 | goat | AF2605-SP | RRID:AB_2108571 |
| KRT13 | rabbit | HPA030877-25UL | RRID:AB_2673641 |
| KRT14 | mouse | 314M-14 | RRID:AB_1159418 |
| KRT5 | rabbit | 305R-14 | RRID:AB_1159459 |
| p63 | mouse | AM418GP | |
| RUNX1 | rabbit | ab92336 | RRID:AB_2049267 |
| UPK2 | rabbit | HPA043312-25UL | RRID:AB_2678420 |
| Alexa Fluor 488 Donkey anti-Goat | Donkey | Life Technologies #A11055 | RRID:AB_2534102 |
| Alexa Fluor 568 Donkey anti-Mouse | Donkey | Life Technologies #A10037 | RRID:AB_2534013 |
| Alexa Fluor 647 Donkey anti-Rabbit | Donkey | Life Technologies #A31573 | RRID:AB_2536183 |
| Alexa Fluor 488 Donkey anti- Rat | Donkey | Life Technologies #A21028 | RRID:AB_2535794 |
| Alexa Fluor 568 Donkey anti-Goat | Donkey | Life Technologies #A11057 | RRID:AB_2534104 |

#### Figure Legends

**Figure S1 related to figure 1. Urothelial markers are present in HCO differentiations. (A)** Schematic of HIO versus HCO differentiation. **(B)** Heatmap of urothelial genes in HIOs compared to HCOs. Heatmap is based on TPM values from RNA-seq data. **(C)** Examples of immunostaining of colon organoids, urothelial organoids, mosaic organoids based on staining for CDH1 (green), CDX2 (red), and KRT13 (white) and counterstained with DAPI .

**Figure S2 related to figure 1. Additional marker(s) expressed in mouse ventral hindgut and urothelium from HCO differentiations. (A)** Feature plots of additional urothelial genes. Wholemount immunofluorescence staining of day an e10.5 mouse embryos for **(B)** CDH1 (blue), CDX2 (red), GATA3 (green), and **(C)** FOXA2 (red), RUNX1 (green).

**Figure S3 related to figure 2 and 3. Gene ontology and transcription factor expression in WNT modulated 10-day hindgut organoids. (A)** Gene ontology analysis of genes that are upregulated in the dorsal versus ventral cloaca. **(B)** Table of most upregulated transcription factors in dorsal cloaca. **(C)** Gene ontology analysis of genes upregulated in day 10 BMP + ICRT3 cultures compared to day 10 BMP + CHIR cultures. **(D)** Table of most upregulated transcription factors in day 10 BMP + ICRT3 cultures compared to day 10 BMP + CHIR cultures. TFs that are common in (**C,D)** are shown in bold.

**Figure S4 related to figure 4. Dorsal ventral patterning by WNT modulation is stable. (A)** Heatmap of human fetal large intestinal transcripts and their expression in BMP + ICRT3 cultures compared to day 35 BMP + CHIR cultures. **(B)** Heatmap of human fetal bladder transcripts and their expression in day 35 BMP + CHIR cultures compared to day 35 BMP + ICRT3 cultures. **(C)** Hypergeometric means tests comparing the transcriptomes of human fetal large intestine with HCOs and human fetal bladder with HUOs.

**Supplementary table 1. List of antibodies used in this study.**
